# False discovery rate estimation and heterobifunctional cross-linkers

**DOI:** 10.1101/239715

**Authors:** Lutz Fischer, Juri Rappsilber

**Author notes:** Corresponding author (JR).

## Abstract

False discovery rate (FDR) estimation is a cornerstone of proteomics that has recently been adapted to cross-linking/mass spectrometry. Here we demonstrate that heterobifunctional cross-linkers, while theoretically different from homobifunctional cross-linkers, need not be considered separately in practice. We develop and then evaluate the impact of applying a correct FDR formula for use of heterobifunctional cross-linkers and conclude that there are minimal practical advantages. Hence a single formula can be applied to data generated from the many different non-cleavable cross-linkers.

## Introduction

Cross-linking mass-spectrometry (CLMS) has become an increasingly popular tool for analyzing protein structures, protein networks and protein dynamics [1–4]. Recently the question of what is the correct error estimation to use with CLMS has been addressed with the help of a target-decoy database approach[5], based on previous work for cross-linked[6,7] and linear peptides[8–11]. This approach to estimating a false discovery rate (FDR) of cross-links is based on the assumption that the cross-linker used is homobifunctional, i.e. have the same reactive group on either end. However, heterobifunctional cross-linkers are also used in the field, for example 1-ethyl-3-(3-dimethylaminopropyl)carbodiimide hydrochloride (EDC)[12–15] or succinimidyl 4,4’-azipentanoate (SDA)[15–21]. It is unclear how far these cross-linker choices affect FDR estimation as they do link different amino acids and consequently one has to consider different search spaces. Here, we provide some theoretical insights on extending the target-decoy approach to FDR estimation when using heterobifunctional cross-linkers, and assess whether it is necessary to use a different formula for FDR estimation. Note that these considerations are for non-cleavable cross-linkers. While MS-cleavable cross-linkers with independent identification of both peptides could be treated the same way, by taking the two identifications as one combined identification, they are currently handled differently for FDR estimation[22,23].

## Results and Discussion

Currently, the most commonly used cross-linkers are non-directional, e.g. when looking at a mass-spectrum of a cross-linked peptide, there is no means to distinguish a cross-link that was formed as peptide A linked to peptide B, than from a cross-link formed as peptide B linked to peptide A. But the most commonly used formula[24–28]

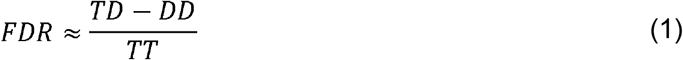

is actually for directional cross-links[5]. Here TT is the number of observed target-target matches (both cross-linked peptides come from the target database), TD is the number of observed target-decoy matches (one linked site comes from the target database and one from the decoy database) and DD stands for the number of decoy-decoy matches (both peptide matches are from the decoy database). A correct formula for the more commonly used non-directional cross-linker (e.g. BS^3^ or DSS) would be[5]:

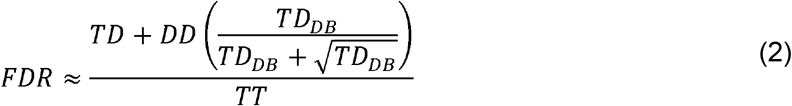

This formula requires knowledge of the number of possible target-decoy pairs in the initial search database (*TD_DB_*). However, the error made by using formula 1 approaches zero relatively fast with increasing database size. Therefore in practical terms the directional formula is also applicable to data of non-directional cross-linkers such as BS^3^ or DSS.

Directionality (or the lack of it) is not the only property of a cross-linker. Cross-linkers can also be homobifunctional or heterobifunctional. For homobifunctional cross-linkers, any peptide in the database that can react with one side of the cross-linker, can also react with the other side. For heterobifunctional cross-linker that is not the case, which has consequences for constructing the target and decoy search space. It leads to distinct databases (set of peptides or residue pairs) for each side of the cross-linker. The formulas used previously, assume a homobifunctional cross-linker.

A set of considerations (see supporting information S1 Derivation of formula) leads us to an FDR estimation formula for non-directional, heterobifunctional cross-linkers:

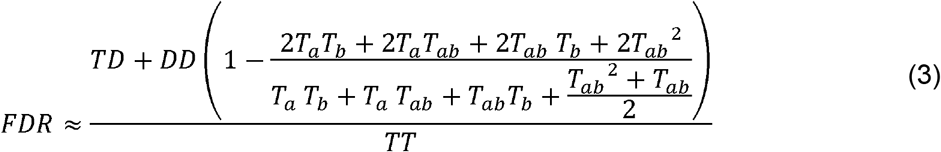

Besides the observed target-target (TT), target-decoy and decoy-target (TD), and decoy-decoy matches, it needs a set of parameters describing the search database (Table 1). As formula 2 can be simplified to formula 1 in all practical terms we wondered how big an error would occur when also using the much simpler formula for directional, homobifunctional cross-linkers (formula 1), in place of formula 3.

**Table 1:** Formula Symbols

| Symbol | Meaning |
| --- | --- |
| $T_a$ | Target entries in the database linkable by side A of the cross-linker |
| $T_b$ | Target entries in the database linkable by side B of the cross-linker |
| $T_{ab}$ | Target entries in the database linkable by both sides the cross-linker |
| $D_a$ | Decoy entries in the database linkable by side A of the cross-linker |
| $D_b$ | Decoy entries in the database linkable by side B of the cross-linker |
| $D_{ab}$ | Decoy entries in the database linkable by both sides the cross-linker |
| TT | Observed target target matches with |
| TD | Observed target decoy and decoy target matches |
| DD | Observed decoy-decoy matches |

The error appears once matches with two decoy peptides are encountered. Before then, one arrives at the same FDR value with formula 3 and 1. Up to this point we have a linear problem (Fig 1a), as we can use the decoys only to model the hits with one wrongly identified partner, and overlook any match to two wrongly identified partners. Statistically, these will be rare, however they are not modeled until a significant number of decoy-decoy matches are encountered.

**Fig 1.**
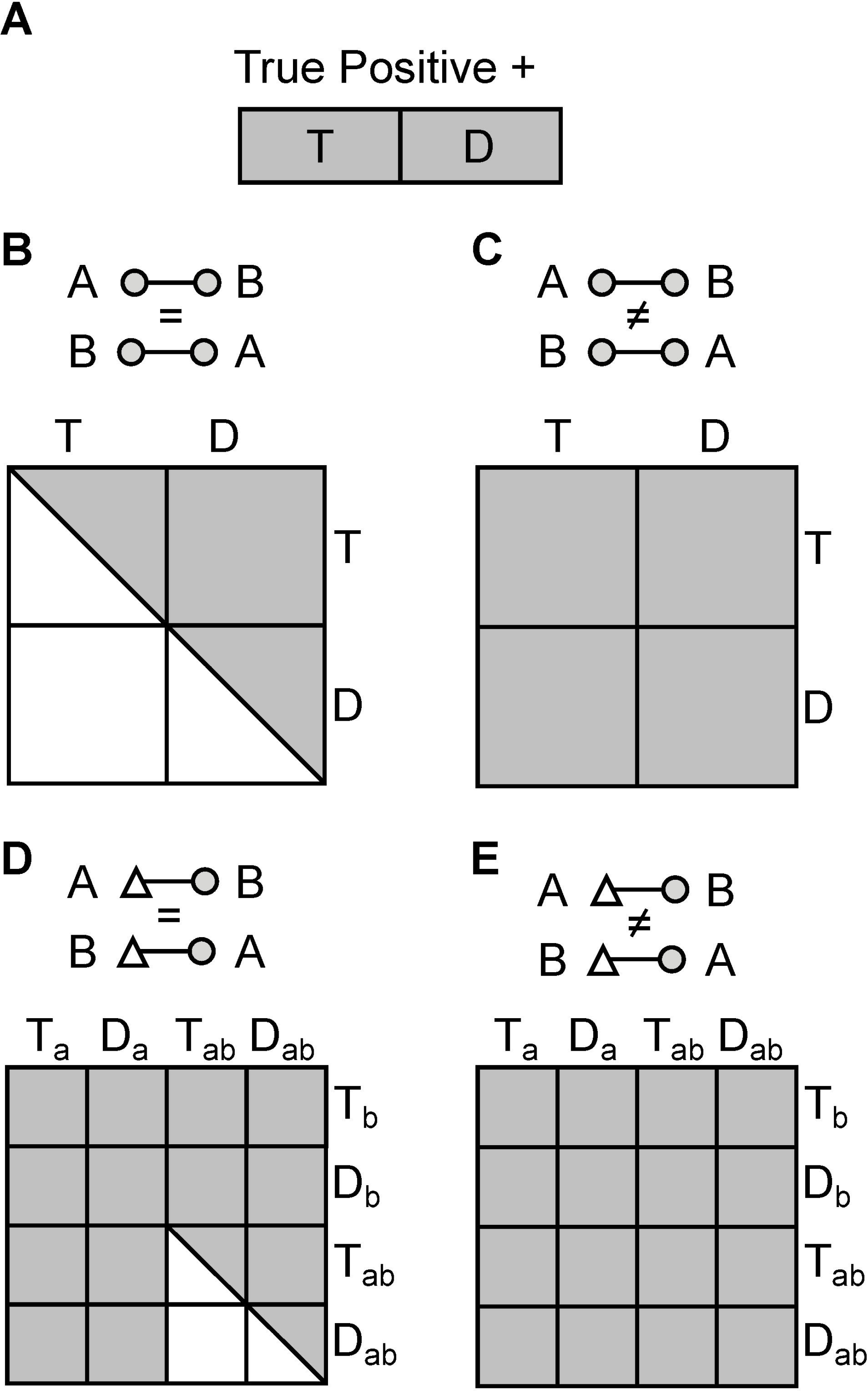
Random search spaces for false positive matches. To model matches where one correct and one incorrect partner are combined requires considering a linear random match space (A). In contrast, when modelling matches with two incorrect partners it requires construction of a quadratic random match space depending on whether the cross-linker is homodimeric, non-directional (B), homodimeric, directional (C), heterodimeric, non-directional (D), or heterodimeric, directional (E).

The situation changes once matches with two decoys are encountered. Here we start modeling how likely we have hits with two wrongly matched partners. The random space for a non-directional heterobifunctional cross-linker is somewhere between the directional and nondirectional spaces for the homobifunctional cross-linker (Fig 1b). In fact the larger the non-overlap is between the two sites of the cross-linker - and therefore the smaller T_ab_ and D_ab_ are - the closer it behaves like a directional, homobifunctional cross-linker and the simplification of formula 1 applies.

The error made when using formula 1 for heterobifunctional cross-linkers is smaller than the error made when using formula 1 for non-directional homo-bifunctional cross-linkers (Fig 2). Already, at 200 entries (i.e. peptide, linkable residues or proteins, depending of what level the FDR should be estimated on [5]) in the database, even for a 100% overlap between both sides of the cross-linker (effectively resulting in a directional homobifunctional cross-linker) the error of FDR estimation incurred by using formula 1 instead of formula 3 should not exceed 1%. For example when cross-linking HSA (585 residues in the active form, of which 129 are Lysine, Serine, Threonine or Tyrosine and the protein amino terminus) with SDA, the maximal error resulting from using formula 1 should be less than 0.2% from the estimated FDR - i.e. 5% would be <5.01% (Table 2). This error is usually smaller than the actual resolution of the FDR estimation[5]. Considering EDC in a second example: there is a 100% non-overlap between both sides of the cross-linker (Lysine, Serine, Threonine, Tyrosine, and the protein amino terminus on one side and Glutamic acid, Aspartic acid, and the protein carboxy terminus on the other side). An FDR calculation using formula 1 would result in the same estimate as using formula 3. At the level of peptides, the situation would look slightly different. Taking HSA cross-linked with EDC and a tryptic digest with four missed cleavages would result in 23 peptides exclusively for one side (T_a_), 31 peptides for the other side (T_b_) and 329 peptides (T_ab_) that could be linked to either side of the cross-linker. This would lead to a maximal error of around 0.45% (i.e. 5% would become 5.023%).

**Fig 2.**
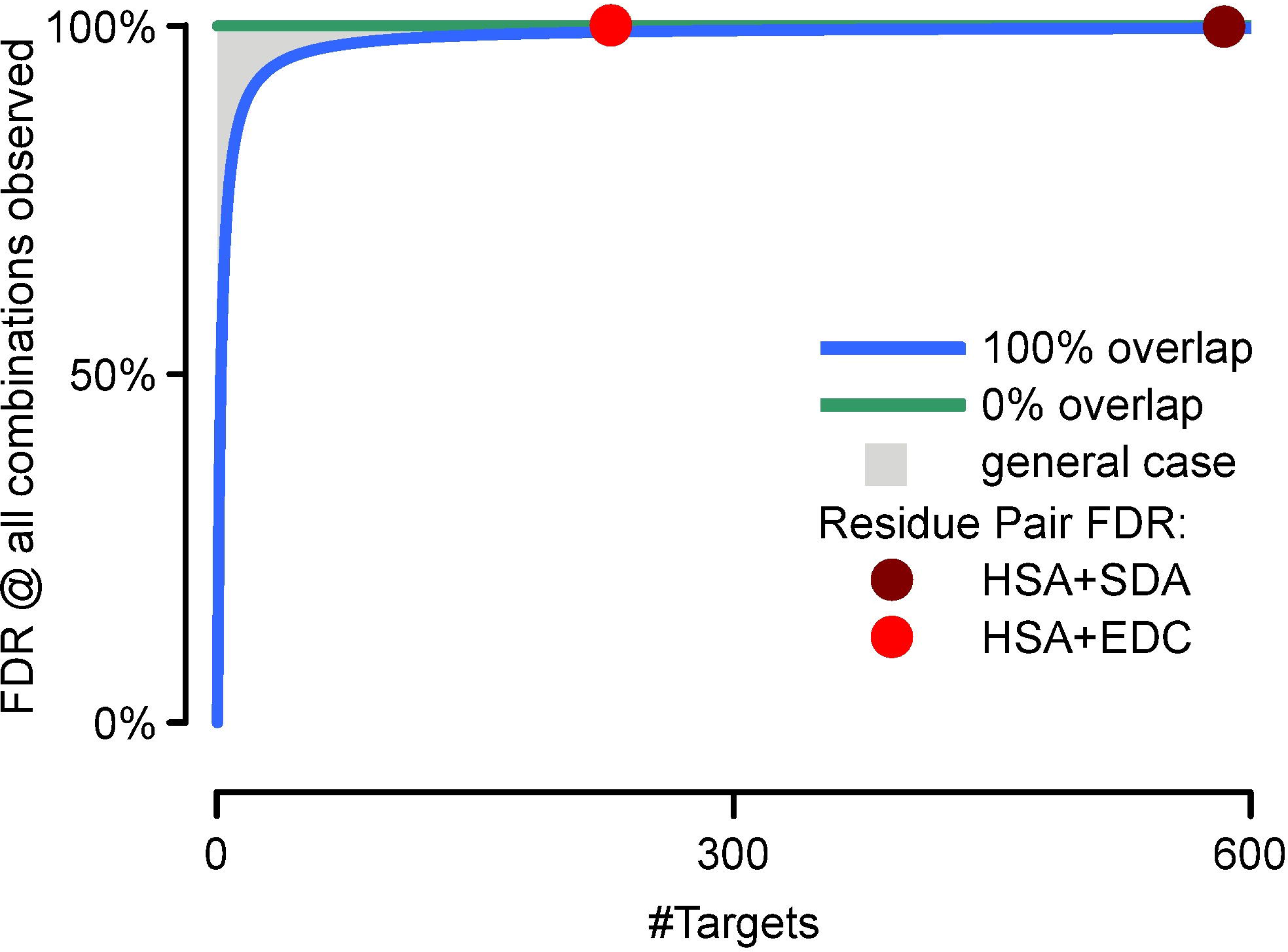
Maximal Error from using formula 1. Maximal expected error when using formula 1, exemplified for the extreme case of every possible combination of links being observed. X-axis is the size of the database and Y-axis is the maximal error. The green and blue line give the border cases of 0% overlap for both sides of the cross-linker and 100% overlap respectively. The gray area represents possible errors for all cross-linker with partial overlap. Residue-level for HSA cross-linked SDA (dark red dot) and HSA cross-linked with EDC (light red dot) are given as reference.

**Table 2:** Examples of maximal expected error when using the simple formula for HSA, cross-linked with either EDC or SDA.

| Cross-Linker | Level | Ta | Tb | Tab | Maximal Error | Formula 1 | Formula 3 |
| --- | --- | --- | --- | --- | --- | --- | --- |
| SDA | *residue pairs* | 0 | 455 | 130 | 0.19% | 5.00% | 5.01% |
|  | *peptide pairs* | 0 | 27 | 360 | 0.48% | 5.00% | 5.02% |
| EDC | *residue pairs* | 99 | 130 | 0 | 0.00% | 5.00% | 5.00% |
|  | *peptide pairs* | 23 | 31 | 329 | 0.45% | 5.00% | 5.02% |

In conclusion, from a theoretical point of view formula 3 is to be used for FDR estimations when working with heterobifunctional cross-linkers. However, for all practical purposes, the simpler formula 1 gives an approximation with an error smaller than the resolution of FDR estimation.

## Supporting information

**S1 File. Derivation of formula**.

